# Degradation of net ecosystem carbon balance in cool-temperate forests by sika deer-induced stand structure alterations and subsequent soil erosion

**DOI:** 10.64898/2026.04.20.719128

**Authors:** Hayato Abe, Dongchuan Fu, Tomonori Kume, Ayumi Katayama

## Abstract

Although natural forests sequester carbon, this function may decline under chronic herbivory by abundant ungulates (hereafter overbrowsing). Specifically, overbrowsing alters stand structure, potentially impair carbon exchanges related to the vegetation. Further, overbrowsing may also accelerate soil erosion, especially in heavy-rainfall regions like Monsoon Asia. We quantified these impacts by estimating net ecosystem carbon balance (NECB; g C m^−2^ yr^-1^) by subtracting heterotrophic respiration (*R*_h_) and lateral carbon export via erosion (*S*_e_) from net primary production (*P*_n_) in southern Kyushu, Japan. Here, about 40-years of overbrowsing by sika deer (*Cervus nippon*) altered mixed broadleaf-conifer stands with presence of understory (PU) into stands with no understory (NU), then further altered into stands dominated by unpalatable shrublands (SR) or stands with canopy gaps (CG). The PU maintained a positive NECB (plot mean = 307.0 g C m^−2^ yr^−1^) because high *P*_n_ (721.9) exceeded the sum of *R*_h_ (175.4) and *S*_e_ (239.5). Alteration from PU into NU converted NECB to negative (−98.2 g C m^−2^ yr^−1^). This was because the suppressed *P*_n_ (400.2 g C m^−2^ yr^−1^) could not offset the sum of *R*_h_ (170.6) and *S*_e_ (327.7). Further degradation into CG caused a profound negative NECB (−894.4 g C m^−2^ yr^−1^), where *P*_n_ (71.9) offset only 7% of the sum of surging *R*_h_ (464.8) and *S*_e_ (501.5). Alteration into SR showed a partially recovered NECB (97.3 g C m^−2^ yr^−1^), driven by shrub growth (*P*_n_; 554.5, *R*_h_; 175.4, *S*_e_; 239.5). However, this recovery is still limited given that lowered shrub biomass and prior topsoil loss via erosion. Our results validate previous findings that stand alteration from PU to SR or CG through NU leads to up to a 49% loss of ecosystem carbon stocks. Preventing stand alteration and soil erosion are key countermeasures against chronic overbrowsing and subsequent erosion.

**Graphical Abstract:** 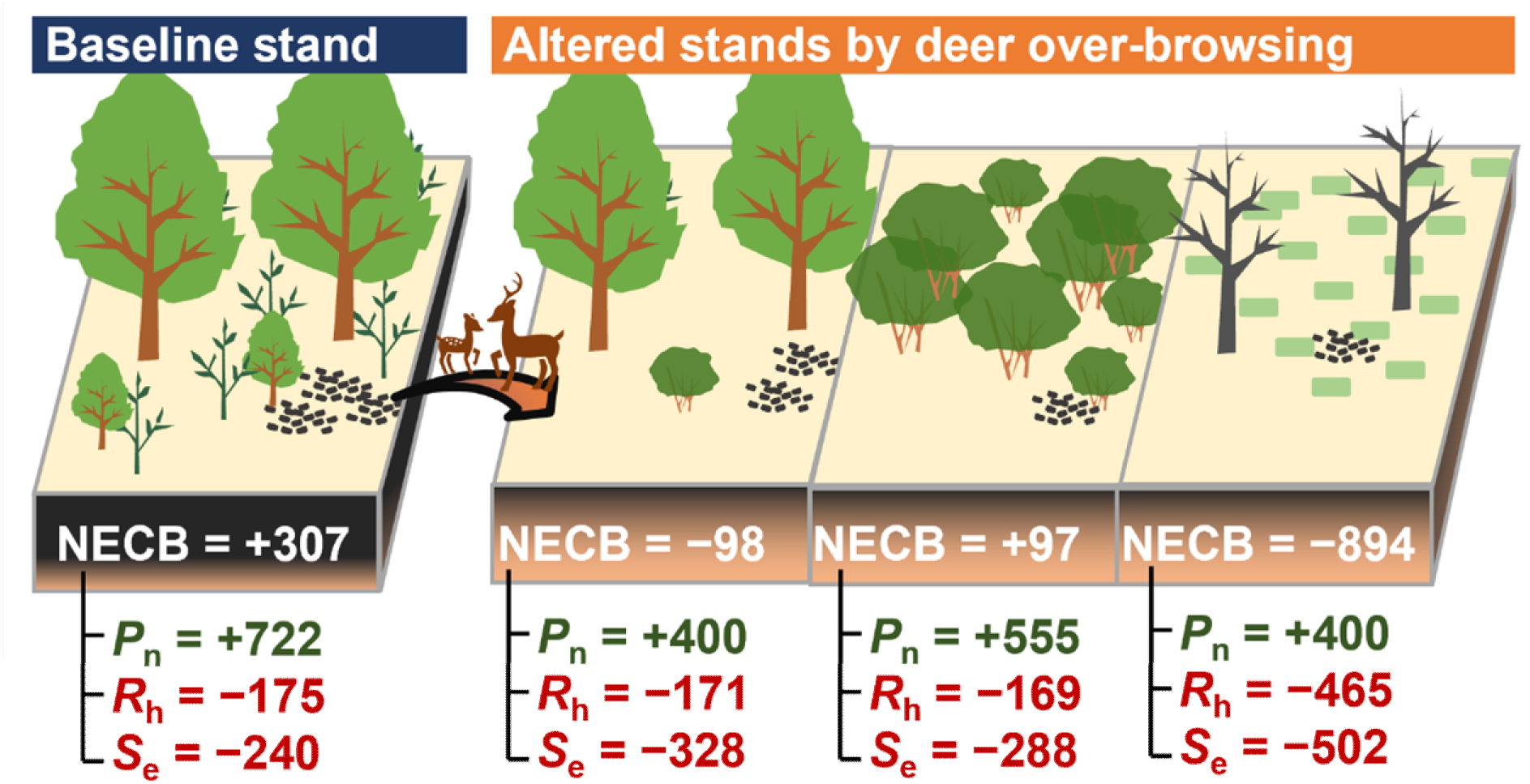

We reported that over 40 years of sika deer overbrowsing and subsequent soil erosion severely degraded the net ecosystem carbon balance (NECB) of mountain forests in Japan. The loss of understory vegetation drove the transition of intact stands into degraded states (no-understory, shrub-dominated, or canopy gaps). Based on field measurements, we quantified that this structural alteration suppressed net primary production (*P*_n_) while increased both heterotrophic respiration (*R*_h_) and lateral carbon loss via soil erosion (*S*_e_). Consequently, the forest shifted from a net carbon sink (+307 g C m⁻² yr⁻¹) to a source (up to −894 g C m⁻² yr⁻¹). These findings provide compelling empirical evidence that increasing ungulate populations, compounded by the rising frequency of heavy rainfall, may severely undermine the carbon sequestration functions traditionally expected of natural forests.

## 1 Introduction

As the largest terrestrial carbon (C) sink, natural forests account for 93% of global forest cover (FAO, 2020). It is believed that as natural forests mature, C sequestration is thought to reach an equilibrium between C uptake and release (Sousa, 1984). Even so, natural forests across the globe have exhibited net C uptake over recent decades (Harris et al., 2021; Pan et al., 2024). This net C uptake is attributed to the forest regrowth from sporadic natural disturbances such as wildfires, windthrow, and pest outbreaks (Harris et al., 2021; Pan et al., 2024; Pugh et al., 2020). During the recovery process, although disturbance-induced tree mortality and C stock losses (e.g., landslide-induced soil loss) instantly export C from ecosystems, high net C uptake is achieved through the growth of recruits and surviving trees (Pugh et al., 2020; Seidl and Turner, 2022). However, there is growing concern that the duration and frequency of natural disturbances are increasing due to changes in human society and global climate change (Harris et al., 2021; Pan et al., 2024). Such frequent and/or prolonged disturbances (hereafter chronic disturbances) may impair the natural forest C sequestration capacity, which has been maintained through the disturbance-regrowth cycle (Seidl and Turner, 2022).

Previous studies have predicted that the impact of chronic disturbances on forest C sequestration may arise through stand structural alterations (McDowell et al., 2008; McDowell and Allen, 2015; Stovall et al., 2019). For example, prolonged droughts are posited to constrain the growth of post-disturbance recruits, shortening the canopy height of the regenerated stand (Allen et al., 2015). Similarly, repeated single disturbances, such as frequent typhoons, can shift tree species composition and tree size distribution (Bonilla-Moheno, 2010; Mitchell, 2013; Ulanova, 2000). Sequential occurrences of multiple disturbance agents also drive similar shifts (Kleinman et al., 2019; Ratajczak et al., 2017; Seidl and Turner, 2022). These stand alterations directly drive the C uptake by vegetation, i.e., net primary production (*P*_n_) (Allen et al., 2015; Pan et al., 2024). Further, these stand alterations may modify C release from dead organic matter, i.e., heterotrophic respiration (*R*_h_), via altered plant-soil feed-backs (Harmon et al., 2011; Williams et al., 2016). In contrast to these prediction, empirical quantification of the extent to which chronic disturbances degrade ecosystem C balance lags behind (Albrich et al., 2020; Dobor et al., 2024; Kleinman et al., 2019; Pan et al., 2024).

Stand alteration may affect not only the gaseous C exchange (i.e., *P*_n_ and *R*_h_), but also non-gaseous lateral C fluxes. In fact, some disturbances, such as ash runoff following wildfires (Flores et al., 2020; Girona-García et al., 2024) and the loss of vegetation and soil organic matter (SOM) due to avalanches (Vacchiano et al., 2015), result in non-gaseous C losses. Under chronic disturbance regimes, such C loss can act repeatedly and cumulatively (Flores et al., 2020). Nevertheless, it remains unclear to what extent the loss of non-gaseous C associated with chronic disturbance affects the dynamics of the NECB. To advance our understanding, there is an urgent need for comprehensive empirical studies that evaluate the effect of chronic disturbances on the NECB through stand alteration by explicitly integrating both gaseous and non-gaseous C pathways.

The concurrent global phenomena of increasing wild ungulate populations and intensifying extreme precipitation regimes represent an emerging chronic disturbance that severely threatens the NECB in natural forests. Recent population increases in wild ungulates have been particularly noticeable in forests across the Northern Hemisphere, reflecting changes in human activity such as the introduction of alien ungulates and the reduction of hunting pressure (Côté et al., 2004; Takatsuki, 2009; Tape et al., 2016). Prolonged high-intensity browsing (hereafter overbrowsing) by ungulates causes widespread declines in understory biomass and species diversity (Suzuki, 2024). This degradation includes the loss of recruitment of canopy species. Thus, species composition and forest structure are altered via regeneration failure in canopy gaps and the dominance of unpalatable plants (Dobor et al., 2024; Suzuki, 2024). Simultaneously, fluctuations in precipitation are increasingly driven by global climate change, raising concerns about the increased frequency and intensity of heavy rainfall events (Zhang et al., 2024). Although heavy rain can trigger water-induced soil erosion, intact understory vegetation usually mitigates this risk. Overbrowsing hinders this function via understory degradation, drastically exacerbating soil erosion. For example, the reduced understory coverage can increase soil erosion rates by up to 33-fold (Mizuno et al., 2021). Since transported SOM is permanently exported from the source area, soil erosion represents a significant lateral non-gaseous C export pathway on hillslopes (Liu et al., 2003).

The mountain forests of Monsoon Asia provide a critical frontline for evaluating this global threat, as they are already experiencing the chronic overbrowsing and subsequent soil erosion, offering profound insights into how such disturbances impact NECB. In southern Kyushu, Japan, overbrowsing by sika deer (*Cervus nippon*) has occurred since 1980s (Saruki et al., 2004). Approximately 40 years of heterogeneous overbrowsing has created a mosaic of different stand alteration within a single landscape (Abe et al., 2024b). Specifically, overbrowsing has triggered the alteration of broadleaf-conifer mixed stands with presence of understory vegetation (presence of understory vegetation; PU, Fig. 1a) into mixed stands with no understory vegetation (no understory vegetation; NU, Fig. 1b). These NU stands have further transitioned into either unpalatable shrublands dominated by Asebi (*Pieris japonica*) (shrublands; SR, Fig. 1c) or persistent canopy gaps (canopy gaps; CG, Fig. 1d) due to the failure of palatable tree regeneration. Simultaneously, under approximately 4000 mm of annual rainfall (Fig. S1), the loss of understory accelerates soil erosion, resulting in tree root exposure of 10–50 cm (Abe et al., 2024c, 2022a). This root exposure hinders tree growth (Abe et al., 2024a), thereby accelerating the transition from NU into SR and CG. Abe et al. (2024b) compared forest C stocks across these stand types and suggested that the transition from PU to SR or CG through NU could lead to up to a 49% loss of total ecosystem C stocks. Building on this, the present study aims to quantify changes in NECB across this overbrowsing-induced degradation gradient (i.e., PU, NU, SR, and CG), with the explicit integration of lateral C export via soil erosion (*S*_e_). Here, we defined NECB as *P*_n_ minus *R*_h_ and *S*_e_, where a positive value indicates net C uptake and a negative value indicates net C release. We expected that stand alteration from PU to SR, and CG via NU would reduce NECB, shifting from positive to negative balance. Further, we predict that the reduction of NECB is driven by a suppression of *P*_n_ coupled with increases in *R*_h_ and *S*_e_. Ultimately, this research provides quantitative basis for understanding the impact of overbrowsing and subsequent soil erosion on the C sequestration potential of natural forest ecosystems.

**Fig. 1.**
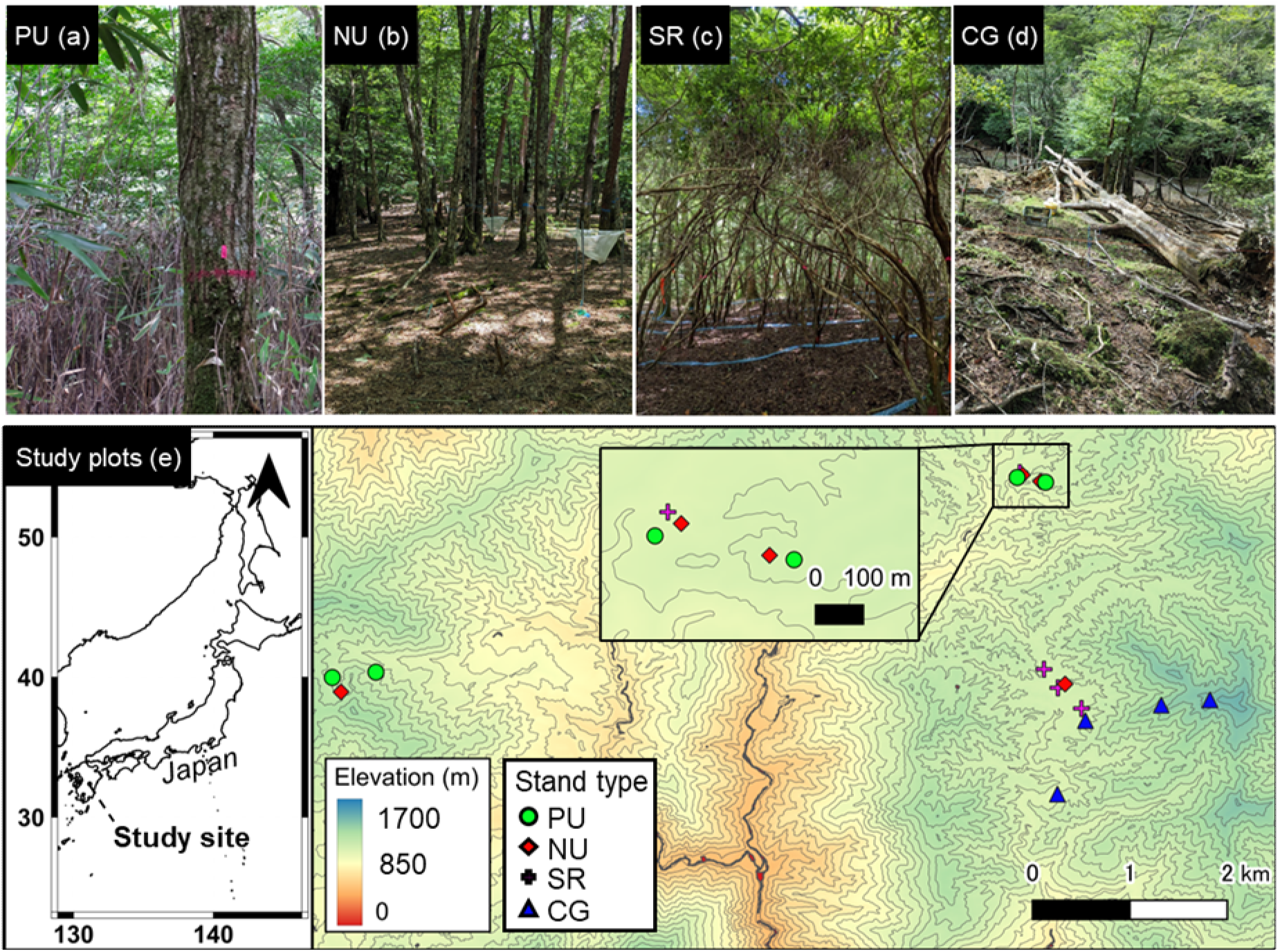
Photographs of (a) broadleaf-conifer mixed stands with presence of understory vegetation (PU), (b) mixed stands with no understory vegetation (NU), (c) stands dominated by shrubs, Asebi (*Pieris japonica*) (SR), and (d) stands with canopy gap areas (CG), as well as (e) location map of study plots. Photographs and the map are adapted from Abe et al. (2024b) under the Creative Commons CC-BY-NC license.

## 2 Methods

### 2.1 Study area

This study was conducted in the Shiiba Research Forest (SRF) of Kyushu University, located in the southern part of Kyushu Island, Japan (32.37274°N, 131.14417°E; 1,031–1,428 m a.s.l.) (Fig. 1e). The mean annual temperature and annual precipitation in 2023 were 12.5°C and 4381.3 mm, respectively (Fig. S1). Prior to the overpopulation of sika deer, the SRF was entirely covered by understory vegetation dominated by the dwarf bamboo (*Sasamorpha borealis*) (Saruki et al., 2004). Following the overbrowsing by deer since the 1980s, understory vegetation cover has decreased across most areas of SRF (Saruki et al., 2004).

In the SRF, 16 survey plots ranging in size from 100 to 400 m^2^ were established in 2022 (Table S1, Fig. 1e), with four plots for each of the four stand types (PU, NU, CG, and SR). Detailed descriptions of these plots are provided in Abe et al. (2024b). Briefly, PU represents the typical stand type of the SRF, where a mixed stand of deciduous broad-leaved trees and evergreen conifers, with seedlings of canopy trees, dwarf bamboo, and other herbaceous species constitute the understory (Fig. 1a). NU represents the mixed stand similar to PU, but lacking understory vegetation due to deer herbivory (Fig. 1b). SR represents the stand formed by the expansion of the unpalatable shrub, Asebi (*P. japonica*) (Fig. 1c). CG represents the stand with canopy gap areas that failed to regenerate palatable trees after overstory decline, characterized by only a few remaining overstory trees and patches of moss forming the understory (Fig. 1d).

### 2.2 Measurements for net ecosystem carbon balance

This study evaluated NECB (g C m^−2^ yr^-1^) by subtracting *R*_h_ and *S*_e_ from *P*_n_ in each plot.

#### 2.2.1 Net primary production

The *P*_n_ was estimated using biometric methods (Ohtsuka et al., 2016), such as the sum of the stand increments of overstory woody biomass (*I*_o_), the stand increments of aboveground understory biomass (*I*_u_), above-ground litter production (*L*), and fine root production (*F*_r_). We measured all components of *P*_n_ in each plot as dry mass basis (g m^−2^ yr^-1^). The dry mass of each component was then converted to C mass (g C m^−2^ yr^-1^) using the C concentration for each component. The C concentrations were obtained from Abe et al. (2024b), except for *I*_o_.

The *I*_o_ was estimated by tracking individual tree growth (Clark et al., 2001). Abe et al. (2024b) recorded the species and stem diameter of all overstory trees with a height >2 m within the entire area of each plot on July 6 to August 8, 2022. We conducted the same measurements on July 10 to 22, 2023. We then estimated the individual biomass of stems, branches, and coarse roots for 2022 and 2023 using allometric equations that required species, stem diameter, and/or species-specific wood gravity (Ichihashi and Katayama, 2024; Ishihara et al., 2015). Allometric equations were shown in Table S2. Species-specific wood gravity data were obtained from Abe et al. (2024b). The biomass increment for each individual tree was calculated by subtracting the biomass in 2022 from the biomass in 2023 for trees surviving in 2023. These data were then aggregated for each plot and divided by the plot area to determine the plot-level biomass increment (g m^−2^ yr^−1^). Finally, these values were converted to C mass (g C m ^−2^ yr^−1^) using a constant C concentration of 0.480 g C g^−1^ (Martin et al., 2018).

The *I*_u_ was estimated as the increment of understory biomass from 2022 to 2023 (Fu et al., 2025). Here, understory vegetation was defined as tree seedlings (height <2 m), dwarf bamboo, mosses, ferns, and other herbaceous plants. Given the absence of understory vegetation in NU and only sparse moss patches in CG (Abe et al., 2024b), *I*_u_ was assumed to be zero for these stand types (Fu et al., 2025). Understory biomass in 2022 was collected in August 1 to 6, 2022, by Abe et al. (2024b). We conducted subsequent evaluations from July 17 to 20, 2023. We established eight subplots (0.5 m * 0.5 m in area) per plot in PU and SR, approximately 1 m outside the plot boundaries on both the left and right flanks. In SR, all understory vegetation within the subplots was harvested and dried in an oven (DKN812, Yamato Scientific, Japan) at 70°C for 1 week. After drying, the samples were pooled for each plot and weighed, calculating dry mass (g m^−2^). In PU, a non-destructive approach was employed to minimize the impact of sampling, as PU are rare in the study area. Instead of harvesting, we recorded the culm density (m^−2^) and height (m) of dwarf bamboo, as well as the coverage (m^2^ m^−2^) and community height (m) of other understory species in each sub-plot. These parameters were converted to dry mass of understory vegetation (g m^−2^) in each sub-plot using established equations (Abe et al., 2025) (Table S2, Fig. S2). Dry mass in each sun-plot was averaged in each pot to obtain representative value. In one SR plot, there was lower the number of newly recruited Asebi seedlings, rather than that outgrew the understory size class (i.e., transitioning from the understory to the overstory compartment). This resulted in a negative value of *I*_u_. For this specific plot, *I*_u_ was set to zero (Table S1).

The *L* was estimated through litterfall collection. We installed four litter traps in each plot in September 2022, with the exception of CG. Due to restricted site access caused by landslides during the study period, only two of the CG plots were equipped with litter traps (Table S1). In PU, SR, and CG, the traps (0.16 m^−2^ in area) were positioned 30 cm above the ground to capture litter from both overstory and understory vegetation. In contrast, traps in NU (1 m^2^ in area) were placed 1 m above the ground to collect overstory litter. The contents were collected approximately every two months from March 2023 to March 2024, and oven-dried at 70°C for 48 hours. Dried samples were pooled for each plot and weighed. For the two CG plots lacking litter traps, *L* was estimated using the average value obtained from the two monitored CG plots (Table S1).

The *F*_r_ was estimated using the ingrowth core method (Katayama et al., 2019; Shimono et al., 2022; Vogt and Persson, 1991). Between October 4 and 6, 2022, soil holes (5 cm in diameter and 30 cm in depth) were created from the center and the four corners of each plot (i.e., 4 stand types * 4 plots * 5 points = 80 sampling points). Ingrowth core bags (5 cm in diameter, 30 cm in depth, 4 mm mesh size) were then inserted into the vacated holes. These bags were filled with root-free soil that had been sieved through a 2-mm mesh. The bags were incubated until the collection period between October 30 and November 4, 2023. After the collection, living fine roots (diameter <2 mm) were extracted from the cores, oven-dried at 70°C for 48 hours, and weighed.

#### 2.2.2 Heterotrophic respiration

The *R*_h_ was estimated as the sum of *R*_h_ from surface soil (*R*_h_SS_) and coarse woody debris (CWD) (*R*_h_CWD_). We estimated *R*_h_SS_ using temperature-driven CO_2_ efflux (μmol CO_2_ m^−2^ s^−1^) equation developed at the SRF (Table S2) (Abe et al., 2025). The equation is based on the CO_2_ efflux from the soil surface, excluding root respiration, as measured using the trench method (Abe et al., 2025). The equation required soil temperature data, then we recorded soil temperature at 30-minute intervals from 2022 to 2023 in one representative plot for each stand type using a thermometer at a depth of 10 cm (S-TMB-M02, Onset, MA, USA). Soil temperature data were then applied to all plots of the corresponding stand type. The annual cumulative CO_2_ efflux was converted to *R*_h_SS_ (g C m^−2^ yr^−1^) based on the molar mass of C.

The *R*_h_CWD_ was estimated using the C stocks of CWD in 2022 (g C m^−2^) and the decay constant (*k*; yr^−1^) (Table S1). The *k* for each plot was estimated using a multiple regression equation (Dai et al., 2021), which incorporates parameters such as mean annual air temperature (°C), annual precipitation (mm), elevation (m a.s.l.), latitude (°), and annual snowfall (kg m^−2^)

(Table S1, S2). The C stock of CWD, elevation, and latitude for each plot were obtained from Abe et al. (2024b). Mean annual air temperature and annual precipitation were derived from 1 km^2^ mesh climate data provided by the Japan Meteorological Agency (https://www.jma.go.jp/jma/en/menu.html). Annual snowfall was assumed to be zero for all plots, as snowfall is negligible in the SRF.

#### 2.2.3 Carbon exports by soil erosion

The *S*_e_ was estimated by multiplying the annual eroded soil depth (*E*; cm yr^−1^) by the C stock of SOM per 1 cm depth (g C m^−2^ cm^−1^). The C stock of SOM per 1 cm depth was derived from data reported by Abe et al. (2024b) for the 0–10 cm depth. The *E* was measured in each plot using the erosion pin method (Gholami et al., 2021; Hart et al., 2017; Kawakami et al., 2026). From October 4 to 6, 2022, we installed 16 pins in each plot (i.e., 4 stand types × 4 plots × 16 pins = 256 pins). Each pin (2.8 cm in diameter, 25 cm in length) was placed at least 1 m apart and inserted approximately 15 cm into the soil. The interface between the pin and the mineral soil surface (excluding the litter layer) was marked. After an incubation period ending between October 30 and November 4, 2023, *E* was measured as the vertical distance from the current mineral soil surface to the initial mark. For five pins buried by upslope sediment (no pins in PU, one pin in NU, two pins in SR, and two pins in CG), *E* was recorded as 0 cm. Additionally, 33 pins that were clearly disturbed by animals (e.g., pulled out or trampled) were excluded from the analysis. Consequently, a total of 223 pins were used for subsequent calculations. We calculated the average value of *E* as plot-level *E* for calculating *S*_e_. In the field, the slope angle (°) at each pin location was also recorded using a clinometer.

### 2.3 Statistical analysis

Using PU as a baseline, we evaluated whether the differences in NECB and its components were statistically significant for each stand type based on plot-level data. These comparisons were performed by a one-sided Dunnet’s test. Specifically, NECB, *P*_n_ and component of *P*_n_ were tested for whether significant decreases relative to PU, while *S*_e_, *R*_h_, and components of *R*_h_ were tested for significant increases.

The amount of *S*_e_ is influenced by variations in slope angle and the C stock of SOM across plots. To examine how the effect of slope angle on *E* varied among stand types, we performed an analysis of covariance (ANCOVA) at the individual pin level, using *E* as the dependent variable, as well as stand type, slope angle, and their interaction as independent variables. Furthermore, we conducted a simple linear regression analysis between the plot-level average of *E* and the C stock of SOM at 0–10 cm depth to assess their relationships. All statistical tests were performed using R software version 4.2.2 (R Core Team, 2024), utilizing the “*stats*” package and the “*multcomp*” package version 1.4–20 (Bretz et al., 2010). The threshold for statistical significance was set at *p* <0.05. For comparisons between NECB and its components, due to the small number of sample plots (four plots) for each stand type, a result with *p* <0.10 was also considered to indicate marginally significant.

## 3 Results

### 3.1 Net ecosystem carbon balance

The net ecosystem carbon balance (NECB) was highest in stands with presence of understory vegetation (i.e., PU, Mean ± SD = 307.0 ± 244.6 g C m^−2^ yr^−1^), followed by stands dominated by Asebi shrubs (i.e., SR, 97.3 ± 158.1), stands with no understories (i.e., NU, −98.2 ± 92.3), and stands with canopy gaps (i.e., CG, −894.4 ± 442) (Table 1, Fig. 2). Compared to PU, the lower NECB was marginally significant for NU (*p* = 0.065) and significant for CG (*p* <0.01). The negative NECB in NU was primarily driven by lower net primary production (*P*_n_). In contrast, NECB in SR was comparable to that in PU, reflecting similar levels of *P*_n_. The negative NECB in CG was attributed to not only lower *P*_n_, but also higher heterotrophic respiration (*R*_h_) and lateral C export via soil erosion (*S*_e_).

**Fig. 2.**
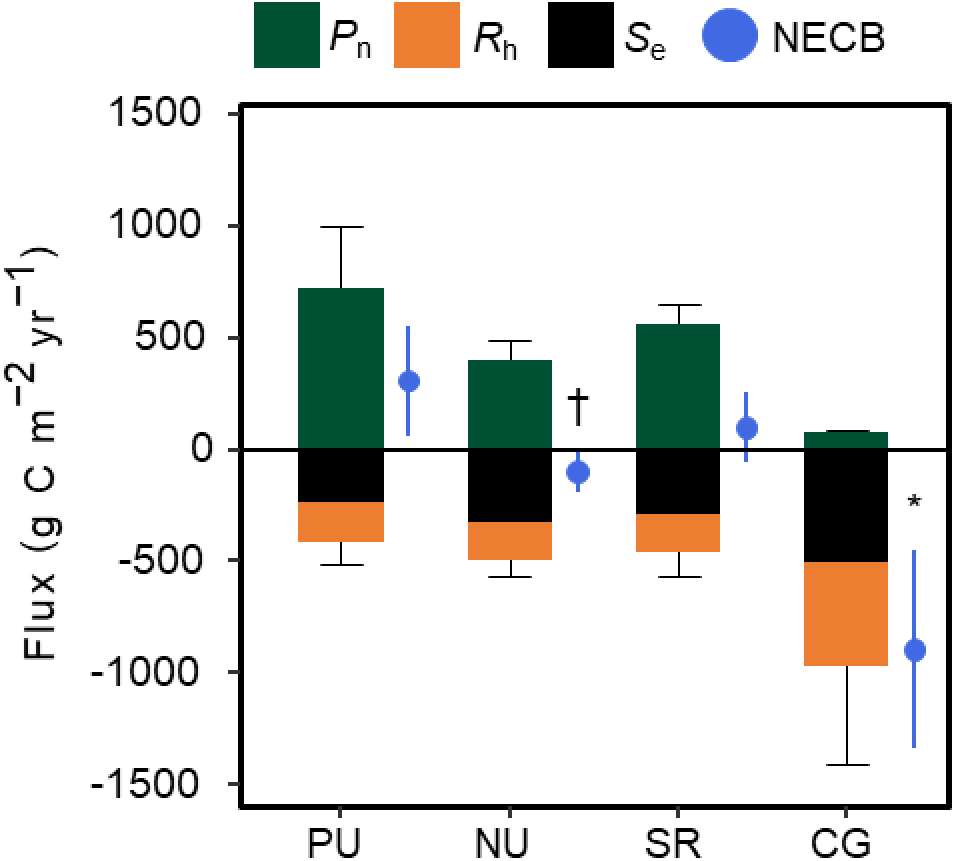
Net ecosystem carbon balance (NECB) of each stand type. The stacking bars indicate the mean value of net primary production (*P*_n_), heterotrophic respiration (*R*_h_), and lateral carbon export via soil erosion (*S*_e_). Upper and lower error bars in stacking bars indicate the SD of *P*_n_ and the sum of *R*_h_ and *S*_e_, respectively. Circles to the right of the stacking bars indicate NECB and its SD. Letters * and † indicate lower NECB from PU with *p* <0.05 and *p* <0.10, respectively.

**Table 1.** Mean (SD) values of net ecosystem carbon balance (NECB, g C m^−2^ yr^−1^) and its components for each stand type. Letters * and † in the carbon flux results indicate the result with *p* <0.05 and *p* <0.10 compared to the PU, respectively.

|  | PU | NU | SR | CG |
| --- | --- | --- | --- | --- |
| NECB | 307.0 (244.6) | -98.2 (92.3) † | 97.3 (158.1) | -894.4 (442)* |
| $P_n$ | 721.9 (277.4) | 400.2 (85.7) * | 554.5 (91.8) | 71.9 (15.3)* |
| $I_o$ | 347.3 (180.9) | 141.2 (63.4) * | 280.6 (97.0) | 9.8 (15.2)* |
| $I_u$ | 70.5 (54.1) | 0<sup>a</sup> | 61.4 (73.1) | 0<sup>a</sup> |
| $L$ | 258.8 (74.8) | 193.2 (13.6) | 177.7 (61.8) | 44.7<sup>b</sup> |
| $F_r$ | 45.3 (20.2) | 65.8 (24.8) | 34.7 (20.6) | 17.5 (10.5) |
| $R_h$ | 175.4 (12.1) | 170.6 (5.0) | 169.2 (95.6) | 464.8 (324.8)* |
| $R_{h\_SS}$ | 148.6 | 128.1 | 109.6 | 136.7 |
| $R_{h\_CWD}$ | 26.8 (12.1) | 42.5 (5.0) | 59.6 (95.6) | 328.1 (324.8)* |
| $S_e$ | 239.5 (97.7) | 327.7 (73) | 287.9 (101.8) | 501.5 (219.4)* |

Abbreviations: $P_n$ ; net primary production, $I_o$ ; stand biomass increment of overstory trees, $I_u$ ; stand biomass increment of understory vegetation, $L$ ; above-ground litter production, $F_r$ ; fine root production, $R_h$ ; heterotrophic respiration, $R_{h\_SS}$ ; $R_h$ of surface soil, $R_{h\_CWD}$ ; $R_h$ of coarse woody debris, and $S_e$ ; lateral carbon export via soil erosion. Superscript a: the $I_u$ in NU and CG was assumed to be zero because these stand types have no understory vegetation or have only small patches of moss species, respectively.
Superscript b: the $L$ of the CG was measured on two plots. Therefore, the SD and statistical differences were not evaluated.

The relative importance of each C flux contributing to the NECB varied across the stand types (Table 1). In PU, *P*_n_ (Mean = 721.9 g C m^−2^ yr^−1^) was the largest flux, followed by *S*_e_ (239.5) and *R*_h_ (175.4). *P*_n_ in PU was approximately 1.7-fold higher than the total C release (i.e., the sum of *R*_h_ and *S*_e_, 414.9 g C m^−2^ yr^−1^). This rank order of fluxes (*P*_n_ > *S*_e_ > *R*_h_) remained consistent in both NU and SR. However, *P*_n_ in NU was lower than the total C release (400.2 vs. 498.3 g C m^−2^ yr^−1^) and *P*_n_ in SR slightly exceeded the total C release (554.5 vs. 457.1 g C m^−2^ yr^−1^). Compared to these three stand types, the rank order of fluxes differed in CG, where *S*_e_ became the dominant flux, followed by *R*_h_ and *P*_n_. In CG, *P*_n_ offset only 7% of the total C release (71.9 vs. 966.3 g C m^−2^ yr^−1^).

### 3.2 Net primary production

The *P*_n_ in NU (400.2 ± 85.7 g C m^−2^ yr^−1^) and CG (71.9 ± 15.3) were 45% and 90% lower than those in PU (721.9 ± 277.4), respectively (Table 1, Fig. S3). These differences were significant (NU; *p* = 0.015, CG; *p* <0.01). In contrast, *P*_n_ in SR remained comparable to that in PU (554.5 ± 91.8 g C m^−2^ yr^−1^, *p* = 0.162). Lower *P*_n_ in NU and CG were primarily attributed to the lower overstory biomass increment (i.e., *I*_o_) (Fig. S3).

The *I*_o_ was highest for PU (347.3 ± 180.9 g C m^−2^ yr^−1^), followed by SR (280.6 ± 97.0), NU (141.2 ± 63.4), and CG (9.8 ± 15.2) (Table 1, Fig. S4). The *I*_o_ in NU and CG was 59% and 97% lower than that in PU, respectively; these differences were significant (NU; *p* = 0.048, CG; *p* = 0.002). The lower *I*_o_ in NU and CG was driven by the lower stem density of palatable trees (Fig. S4). In contrast, SR showed the high stem density of Asebi with stem diameter <10 cm, sustaining high *I*_o,_ which was comparable to that of PU (*p* = 0.718) (Fig. S4).

No significant differences were observed in above-ground litter production (*L*) among PU (258.8 ± 74.8 g C m^−2^ yr^−1^), NU (193.2 ± 13.6 g C m^−2^ yr^−1^, *p* = 0.227), and SR (177.7 ± 61.8 g C m^−2^ yr^−1^, *p* = 0.127) (Table 1). The *L* in CG showed lower value compared to PU (44.7 g C m^−2^ yr^−1^), while SD and *p*-values were not calculated for this stand type due to the limited sample size. Further, no differences in fine root production (*F*_r_) were observed among PU (45.3 ± 20.2 g C m^−2^ yr^−1^), NU (65.8 ± 24.8, *p* = 0.361), SR (34.7 ± 20.6, *p* = 0.792), and CG (17.5 ± 10.5, *p* = 0.163) (Table 1).

### 3.3 Heterotrophic respiration

Total amount of *R*_h_ in NU (170.6 ± 5.0 g C m^−2^ yr^−1^) and SR (169.2 ± 95.6) was comparable to that in PU (175.4 ± 12.1) (NU; *p* = 0.764, SR; *p* = 0.768) (Table 1, Fig. S5). In contrast, *R*_h_ in CG (464.8 ± 324.8 g C m^−2^ yr^−1^) was significantly 2.6-fold higher than that in PU (*p* = 0.04). These differences were primarily driven by the high amount of *R*_h_ from coarse woody debris (CWD) (i.e., *R*_h_CWD_). The *R*_h_CWD_ in CG was 328.1 ± 324.8 g C m^−2^ yr^−1^, about 12-fold higher than that in PU (26.8 ± 12.1 g C m^−2^ yr^−1^, *p* = 0.033) (Table 1). *R*_h_ from surface soil (*R*_h_SS_) was comparable among stand types: 148.6, 128.1 109.6, and 136.7 g C m^−2^ yr^−1^ for PU, NU, SR, and CG, respectively (Table 1). The seasonal variations in soil temperature used to estimate *R*_h_SS_ also did not differ strongly among stand types (Fig. S6).

### 3.4 Lateral carbon exports via soil erosion

The *S*_e_ in NU (327.7 ± 73.0 g C m^−2^ yr^−1^) and SR (287.9 ± 101.8) were comparable to that in PU (239.5 ± 97.7) (NU; *p* = 0.366, SR; *p* = 0.544) (Table 1). The *S*_e_ in CG (501.5 ± 219.4 g C m^−2^ yr^−1^) was significantly about 2-fold higher than that in PU (*p* = 0.023). At the individual pin level, the annual eroded soil depth (*E*) was positively related to slope angle across all stand types (Fig. 3). The result of ANCOVA revealed that slope angle (*p* <0.01) and stand type (*p* <0.01) had significant effects on *E*, while the interaction between stand type and slope angle was not significant (*p* = 0.16) (Fig. 3). Specifically, the interception of the regression line (i.e., *E* at a 0° slope angle) were higher in NU, SR, and CG than in PU (Fig. S7). Furthermore, a significant negative relationships was observed between the plot-level average of *E* and the C stock of SOM at 0–10 cm depth (Fig. 4). This means higher *E* was offset by lower SOM in NU and SR; consequently, *S*_e_ in these stand types did not show a significant difference from PU.

**Fig. 3.**
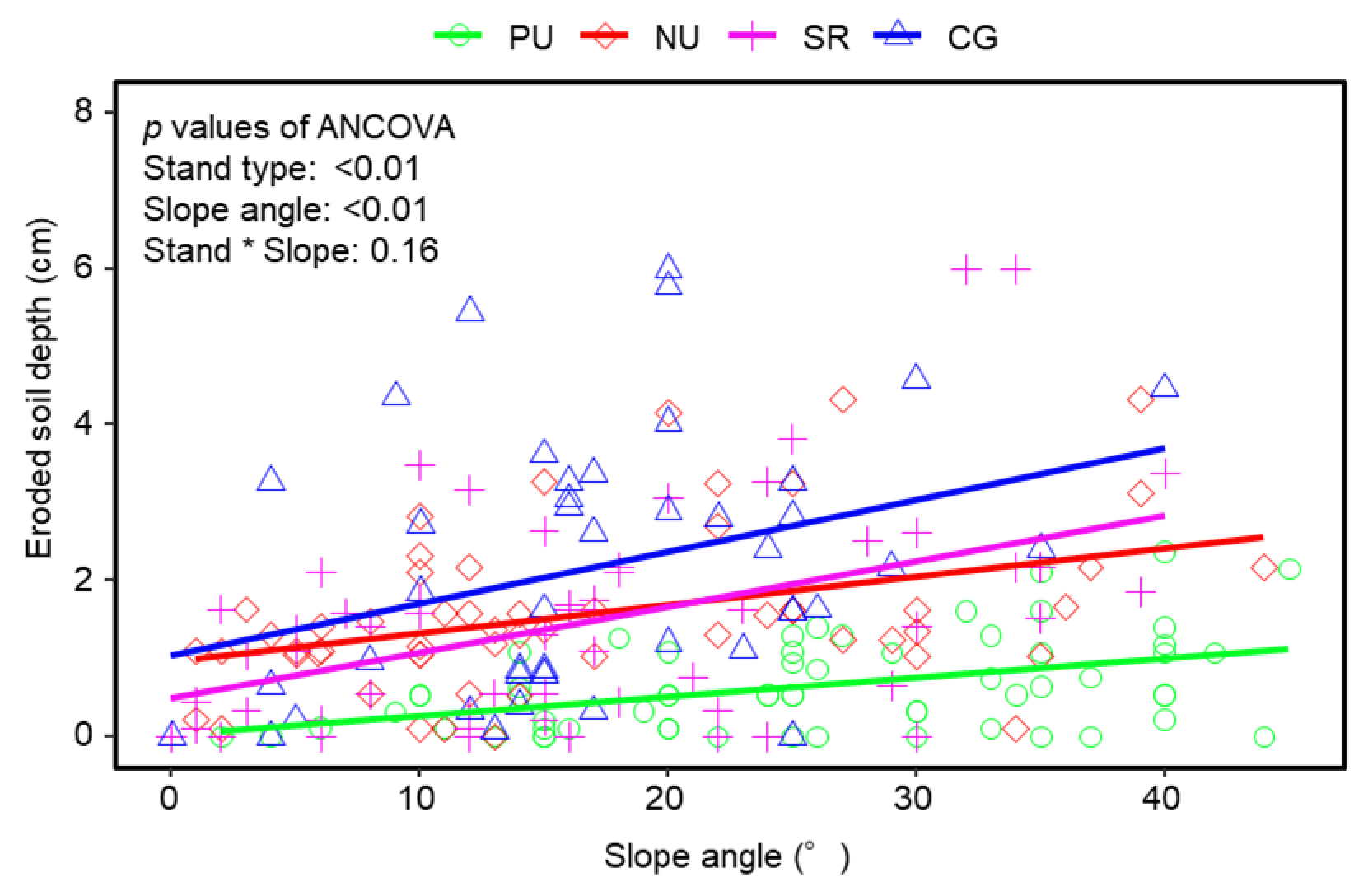
Effect of slope angle on eroded soil depth (*E*) in each stand type. Dots and line refer to the result of each pin and regression line in each stand type.

**Fig. 4.**
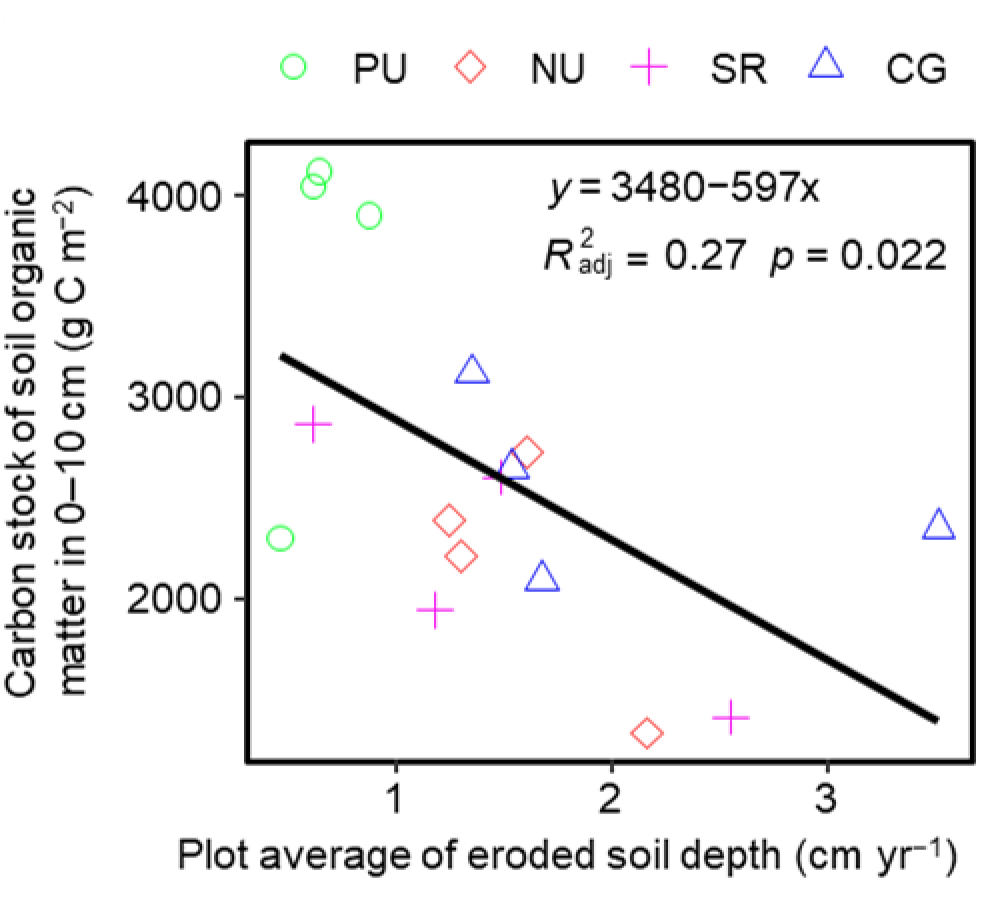
Relationships between carbon stock of soil organic matter at 0–10 cm depth and plot average of eroded soil depth (*E*). The black line indicates the regression line.

## 4 Discussion

### 4.1 Impacts of overbrowsing and soil erosion on net ecosystem carbon balance

Our study quantified the effect of chronic overbrowsing and soil erosion on NECB. Assuming that overbrowsing triggers a successional shift from PU to SR or CG via NU, the NECB was reduced from 307 to −894 g C m^−2^ yr^−1^ (Fig. 5). Specifically, alteration from PU to NU reshaped NECB to a net C source. This degradation was further exacerbated in CG, which exhibited the most severe net C source. In contrast, the colonization of unpalatable shrubs in SR partially restored net C uptake. Nevertheless, its C uptake capacity remained substantially lower than that of PU. These declines in NECB supported the predictions of previous study that ecosystem C stocks gradually decrease as PU alters into SR or CG via NU (Abe et al., 2024b) (Fig. 5). Overall, our results demonstrate that chronic overbrowsing and subsequent soil erosion severely compromises the C sequestration potential of monsoon mountain forests.

**Fig. 5.**
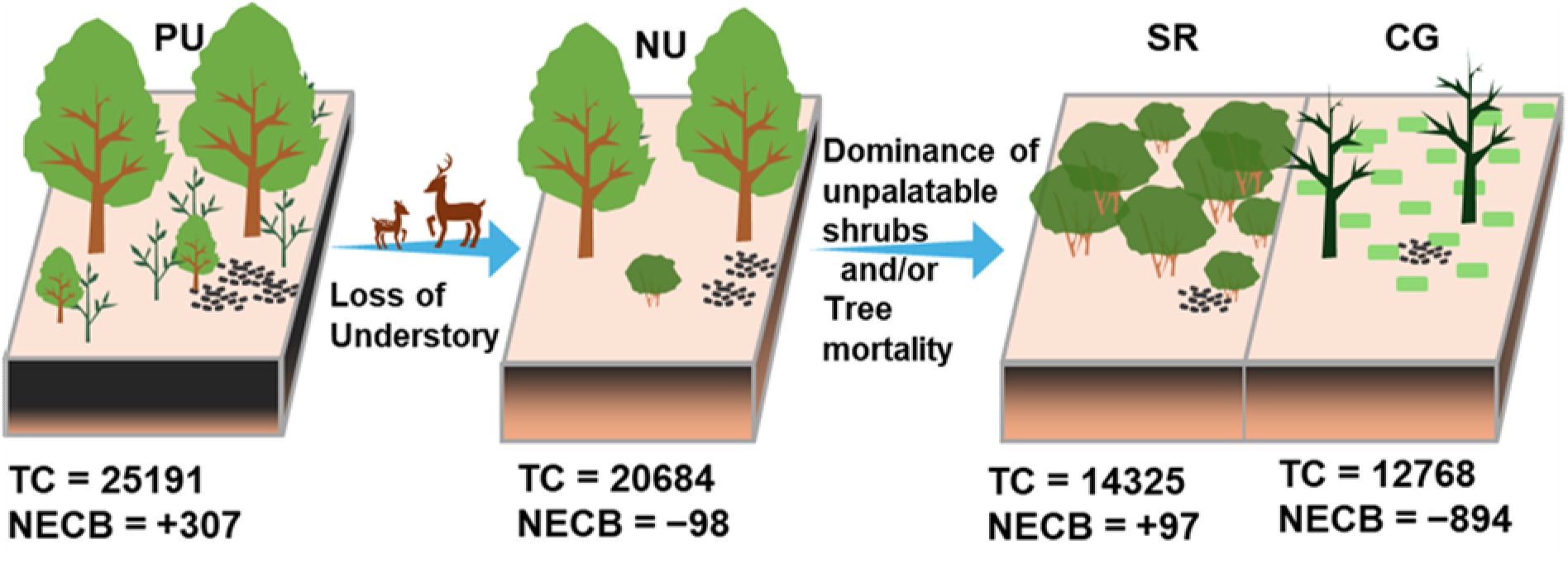
Conceptual diagram of stand structural alterations with total carbon stock (TC, g C m^−2^) and net ecosystem carbon balance (NECB, g C m^−2^ yr^−1^) due to sika deer overbrowsing and subsequent soil erosion. Values represent the mean value of each stand type.

In addition to the absolute changes in NECB, our results revealed shifts in the relative importance of NECB components (i.e., *P*_n_, *R*_h_, and *S*_e_) (Table 1, Fig. 2). In PU, NU, and SR, the ecosystem maintained a *P*_n_-dominated state (*P*_n_ > *S*_e_ > *R*_h_). The transition from a C sink in PU to a C source in NU was not driven by a reordering of this state, but rather by a suppression in the amount of *P*_n_, which fell below the total C release (*R*_h_ + *S*_e_). Conversely, the severe C loss observed in CG, characterized by an export-dominated state (i.e., *S*_e_ > *R*_h_ > *P*_n_). These changes in C flux were caused by a decrease in *P*_n_ as well as an increase in *R*_h_ and *S*_e_ resulting from stand alteration.

The amount of *S*_e_ exceeded *R*_h_ in all stand types (Table 1), identifying it as a dominant C export pathway in our study areas. Although *S*_e_ is often neglected in conventional C flux assessments (Noske et al., 2024), the critical role of *S*_e_ has been pointed out in specific contexts, such as wildfires (Girona-García et al., 2024; Noske et al., 2024) and/or high-precipitation ecosystems like the tropical Amazon (Flores et al., 2020). Our findings suggest that, similar to these cases, integrating *S*_e_ into NECB is essential to accurately capture the shift from C sink to source in monsoon Asian ecosystems under overbrowsing. Given that climate change is projected to increase the frequency and intensity of heavy rainfall globally (Zhang et al., 2024), this integration is imperative not only for current high-precipitation regions like Monsoon Asia, but universally for mountainous ecosystems where extreme precipitation events are anticipated to intensify in the future.

Compared to sporadic disturbances, the impact of chronic overbrowsing and soil erosion on the NECB appears exceptionally severe due to its persistent nature. Typically, sporadic disturbances initially cause a reduction in NECB (Liu et al., 2011). Then, within a few decades, the NECB usually rebounds, sometimes exceeding pre-disturbance uptake levels. For example, Murayama et al. (2024) used the eddy covariance method to monitor the net ecosystem production (NEP, equivalent to *P*_n_ minus *R*_h_) of a Japanese cool-temperate forest for approximately 25 years. They found that a typhoon initially reduced the NEP by 67% (from 289 to 96 g C m^−2^ yr^−1^). This decline was mitigated within two years (225 g C m^−2^ yr^−1^), and by eight years post-disturbance, NEP had increased by 54% (444 g C m^−2^ yr^−1^) compared to the pre-event baseline. Similar dynamics have been reported for wildfires (Vargas et al., 2008), landslides (Schomakers et al., 2019), and insect outbreaks (Hansen, 2014; Morehouse et al., 2008). This recovery occurs because such disturbances, while temporarily reducing C stocks, often stimulate the growth of surviving trees and facilitate recruitment (Hansen, 2014; Morehouse et al., 2008; Murayama et al., 2024; Schomakers et al., 2019; Vargas et al., 2008). In contrast, chronic overbrowsing and soil erosion drive a more persistent degradation, shifting forests into a prolonged net C source. This fundamental difference implies that the reductions in NECB cannot be fully compensated by time alone, suggesting that current global terrestrial carbon models—which largely overlook chronic animal-driven structural changes and non-gaseous C fluxes—may significantly overestimate the long-term resilience and sequestration capacity of natural forests.

### 4.2 Possible processes reducing net ecosystem carbon balance

The *P*_n_ was reduced in NU and CG, but not in SR. In NU and CG, the reduction in *P*_n_ was primarily driven by *I*_o_ (Table 1, Fig. 2). We attribute the decrease in *I*_o_ to two key mechanisms. First, the individual growth of the overstory trees is likely impaired by soil erosion. As documented by Abe et al. (2024a, 2024b), the loss of understory vegetation exposes tree roots by soil detachment, hindering tree growth via water uptake limitations. Second, loss of understory vegetation suppresses the recruitment of palatable canopy species, leading to a long-term reduction in stem density (Enoki et al., 2017) (Fig. S4). In SR, *P*_n_ appeared to recover by the growth of Asebi. However, this recovery is limited. Since Asebi is a shrub species that cannot attain the stature of canopy trees (Ichihashi and Katayama, 2024)—which typically reach several dozen meters—these shrublands will not achieve the C storage capacity of PU.

An increase in *R*_h_ was observed in CG compared to PU (Table 1, Fig. S5). This result was primarily driven by the *R*_h_CWD_. In the present study, *R*_h_CWD_ is determined by the product of the decay constant (*k*) and the C stock of the CWD. The estimated *k* remained relatively consistent across stand types (ranging from 0.105 to 0.121, Table S1), indicating that the variance in *R*_h_CWD_ was almost dictated by the CWD stocks. Previous studies that directly measured CO_2_ fluxes from CWD (Abe et al., 2022b; Gora et al., 2019; Jomura et al., 2007) also indicate that the variability in *R*_h_CWD_ is almost explained by CWD stock rather than CO_2_ efflux rate from CWD. These results highlight that overstory mortality in CG not only degrade *P*_n_ but also creates a long-term C source through the gradual decomposition of accumulated CWD.

Compared to PU, an increase in *S*_e_ was not observed in the NU and SR, but was confirmed only in the CG (Table 1). Nevertheless, we believe that the shift from PU to NU and SR increased *S*_e_, as in the case of CG. This is because *S*_e_ in the present study does not evaluate the total C loss since overbrowsing began, but rather is a snapshot result at a somewhat advanced stage. The present study evaluated *S*_e_ by multiplying the erosion rate (i.e., *E*) by the C stocks of SOM at surface layer. Although NU and SR showed increased *E* (Fig. 3), the C stocks of SOM in these stands were lower than that in PU (Fig. 4). This means that the increased *E* was offset by reduced SOM, sustaining *S*_e_ comparable to that in PU. Similarly, in CG, there was an increase in *E* and a decrease in SOM; however, the increase in *E* exceeded the decrease in SOM, resulting in a higher *S*_e_ level than in the PU. These results indicate that the seemingly comparable *S*_e_ in NU and SR to that in PU does not imply a low impact of erosion; rather, it is a direct consequence of the severe depletion of topsoil SOM caused by chronic erosion prior to our measurements (Abe et al., 2024b).

The fate of the eroded SOM remains a subject of ongoing debate. Some studies suggest that eroded SOM can act as a C sink through deposition and subsequent burial (Van Oost and Six, 2023). In the present study area, much portion of the eroded soil is likely exported to aquatic systems rather than being redistributed within the terrestrial landscape (Chiwa, 2021). Chiwa (2021) reported increased water turbidity and sediment-associated nutrient loading in streams following deer over-populations, suggesting that overbrowsing-induced erosion facilitates a direct lateral export of C from the forest ecosystem to rivers.

### 4.3 Implications for forest management and limitations

Our findings offer insights into mountain forest management in Monsoon Asia and other regions facing similar chronic disturbances. To combat overbrowsing-induced forest degradation, previous management strategies have primarily focused on preventing direct vegetation consumption by ungulates. Consequently, significant attention has been given to population control measures, such as hunting (Iijima et al., 2023), alongside mitigation efforts like the installation of ungulate exclusion fences (Iijima, 2026). From the perspective of preserving forest C sequestration, our results support the continued effectiveness of these conventional interventions. Additionally, our study highlights that localized measures to prevent soil erosion are equally as critical as these traditional approaches. Specifically, proactive interventions such as the installation of wooden steps and ground netting (Sun et al., 2020) are vital to prevent the permanent loss of SOM. Therefore, these localized erosion-control measures must be actively implemented with traditional countermeasures to comprehensively safeguard the ecosystem.

Implementing these countermeasures across an entire forest landscape is logistically challenging. Hence, a critical next step is to maximize the cost-effectiveness of landscape conservation efforts by tailoring specific interventions to the appropriate stage of stand degradation. Here, classifying types of stand alteration is useful to perform triage. For instance, deploying ungulate exclusion fences should be highly prioritized in NU to promote rapid understory recovery before severe soil erosion begins. Furthermore, in late-stage NU and CG, where immediate vegetation recovery cannot be expected, physical erosion control measures must be prioritized. Integrating remote sensing technologies would allow for the continuous monitoring of these stand structural alterations over large areas, facilitating the identification of critical zones and ensuring the optimal deployment of matching countermeasures (Uden et al., 2019).

In order to make such forest management more robust, several spatiotemporal limitations inherent in the present study must be addressed. Spatially, our observations were restricted to relatively small plot sizes within a specific Monsoon Asian climate characterized by high precipitation, which inherently exacerbates soil erosion. This site-specific context limits the immediate generalizability of our findings. Temporally, the single-year measurement period may not fully capture the interannual variability of the NECB and its constituent components (Murayama et al., 2024). To enhance the universality of these findings, future research should prioritize long-term monitoring and cross-site comparisons under diverse meteorological, topographical, and disturbance regimes. These future efforts will provide essential tools for accurately assessing and forecasting the consequences of chronic overbrowsing and soil erosion.

## 5 Conclusion

The present study demonstrates that stand alterations from PU to SR or CG via NU under chronic sika deer overbrowsing and subsequent soil erosion lead to a reduction in the NECB, ultimately resulting in the loss of ecosystem C stocks reported by Abe et al. (2024b). The alteration from PU to NU, characterized by the loss of understory vegetation, triggers an increment of *S*_e_. This accelerated erosion, in turn, induces a decline in tree growth, which, combined with overstory mortality from other natural disturbances (e.g., typhoons), competition, and/or senescence, causes a reduction in *P*_n_. Consequently, the NECB in NU shifts to a negative value, transforming the forest into a net C source. As this condition persists, the combination of overstory mortality and recruitment failure drives the alteration from NU to CG, which, accompanied by an increase in *R*_h_CWD_, will further exacerbate the negative NECB. In cases where unpalatable shrubs colonizes these degraded areas, the stand alters to SR. While SR exhibits a recovery of NECB to positive values due to the growth of unpalatable shrubs, the overall recovery of C stocks is likely limited. This is because C in topsoil may already be lost through erosion prior to shrub colonization, and the C sequestration potential of unpalatable shrubs is lower than that of other overstory trees. Our findings motivate forest managers and governors to consider such chronic disturbances may lead to a permanent decline in forest C sequestration, and to develop countermeasure packages for it.

## Abbreviations and acronyms

C: Carbon
CG: Stands with canopy gap areas
CWD: Coarse woody debris
*E*: Annual eroded soil depth
*I*_o_: Stand-scale overstory biomass increment
*I*_u_: Stand-scale understory biomass increment
*k*: Decay constant
*L*: Above-ground litter production
NECB: Net ecosystem carbon balance
NU: Mixed stand with no understory vegetation
*P*_n_: Net primary production
PU: Mixed stand of broadleaf and conifer trees with presence of understory vegetation
*F*_r_: Fine root production
*R*_h_: Heterotrophic respiration
*R*_h_CWD_: Heterotrophic respiration from coarse woody debris
*R*_h_SS_: Heterotrophic respiration from surface soil
*S*_e_: Carbon export by soil erosion
SOM: Soil organic matter
SR: Shrubland stands dominated by the Asebi (*Pieris japonica*)
SRF: Kyushu University Shiiba Research Forest.

## Declarations

### Availability of data and material

The plot-level data of NECB components is available in Table S1. The plot-level data of C stocks and stand structural data can be found in supplementally materials in Abe et al. (2024b). More detailed data will be made available upon request.

### Authors’ contributions

**Hayato Abe:** Conceptualization, Methodology, Data curation, Investigation, Formal analysis, Funding acquisition, Visualization, Project administration, Writing – original draft. **Dongchuan Fu:** Writing - review & editing, Investigation, Funding acquisition. **Tomonori Kume:** Writing - review & editing, Supervision, Investigation, Funding acquisition. **Ayumi Katayama:**

Writing - review & editing, Investigation, Conceptualization, Methodology, Funding acquisition

## Acknowledgements

We deeply appreciate Mr. Zhouqiang Li for his significant contributions to data collection. We also appreciate Dr. Tadamichi Sato, Ms. Mimori Oyamada, and Mr. Edwin Mukanza Imfumu for their support in field investigation. We acknowledge technical staff of the Shiiba Research Forest of Kyushu University for their help and support in data acquisition in the field. The English grammar of this manuscript were partially refined using Gemini 1.5 Flash (URL: https://gemini.google.com/), a generative AI tool. The authors have reviewed and edited the output to ensure accuracy and take full responsibility for the content of the final manuscript.

