## Supplemental materials for "Degradation of net ecosystem carbon balance in cool-temperate forests by sika deer-induced stand structure alterations and subsequent soil erosion"

# in

Hayato Abe: 0000-0001-6092-0130

Donchuan Fu: 0009-0006-8928-8523

Tomonori Kume: 0000-0001-6569-139X

Ayumi Katayama: 0000-0002-4200-8575

**Table S1.** Plot-level raw data (continue to next page)

| Stand type | Plot rep. | Latitude | Longitude | Plot shape (m) | Plot area ($m^2$) | Slope angle (°) | Elevation (m) | Stand age (yr) | Stand height (m) | Stem density ($ha^{-1}$) | Mean diameter (cm) |
|---|---|---|---|---|---|---|---|---|---|---|---|
| PU | A | 32.398934 | 131.1714 | 20 * 20 | 400 | 7.9 | 1031 | 104 | 18.3 | 575 | 26.1 |
| PU | B | 32.398406 | 131.1745 | 15 * 15 | 225 | 21.1 | 1059 | 102 | 15.6 | 8711 | 2.8 |
| PU | C | 32.377135 | 131.0967 | 15 * 10 | 150 | 20.1 | 1131 | 106 | 14.2 | 1867 | 14.3 |
| PU | D | 32.377713 | 131.1015 | 15 * 15 | 225 | 37.3 | 1967 | 92 | 14.2 | 2489 | 10 |
| NU | A | 32.399208 | 131.172 | 20 * 20 | 400 | 10.3 | 1028 | 104 | 20.2 | 425 | 25.4 |
| NU | B | 32.398503 | 131.174 | 20 * 20 | 400 | 26.8 | 1065 | 142 | 16 | 2625 | 7.7 |
| NU | C | 32.376424 | 131.1767 | 20 * 20 | 400 | 11.2 | 1089 | 108 | 16.8 | 700 | 24.6 |
| NU | D | 32.375555 | 131.0977 | 20 * 20 | 400 | 25.1 | 1151 | 97 | 14 | 2375 | 11.9 |
| SR | A | 32.399465 | 131.1717 | 15 * 15 | 225 | 7.5 | 1024 | 43 | 4.8 | 12444 | 5.9 |
| SR | B | 32.378033 | 131.1744 | 15 * 15 | 225 | 19.1 | 1071 | 47 | 5.1 | 9867 | 3.9 |
| SR | C | 32.376005 | 131.1759 | 15 * 10 | 150 | 6.6 | 1063 | 71 | 3.6 | 19067 | 4.7 |
| SR | D | 32.373761 | 131.1785 | 10 * 10 | 100 | 32 | 1080 | 39 | 2.8 | 21100 | 3.2 |
| CG | A | 32.3746989 | 131.1925 | 20 * 10 | 200 | 15.1 | 1428 | – | – | 150 | 29.4 |
| CG | B | 32.3741265 | 131.1872 | 20 * 10 | 200 | 21.8 | 1400 | – | – | 100 | 38.4 |
| CG | C | 32.364427 | 131.1758 | 7 * 15 | 105 | 28.9 | 1045 | – | – | 95 | 1.1 |
| CG | D | 32.372434 | 131.1789 | 15 * 8 | 120 | 13.6 | 1103 | – | – | 83 | 2.2 |

Note: Stand age, stand height, stand density, and mean diameter are obtained from Abe et al. (2024):

Abe, H., Kume, T., Katayama, A., 2024. Reduction in forest carbon stocks by sika deer-induced stand structural alterations. For. Ecol. Manage. 562, 121938. https://doi.org/10.1016/j.foreco.2024.121938

**Table S1 (Continued).**

| Stand type | Plot rep. | Carbon fluxes (g C $m^{-2}$ $yr^{-1}$) | | | | | | | | | | $E$ (cm $yr^{-1}$) | SOM at 0-10 cm (g C $m^{-2}$) | $k$ ($yr^{-1}$) |
|---|---|---|---|---|---|---|---|---|---|---|---|---|---|---|
| | | NECB | $P_n$ | $S_e$ | $R_h$ | $I_o$ | $I_u$ | $L$ | $F_r$ | $R_{h\_SS}$ | $R_{h\_CWD}$ | | | |
| PU | A | 438.4 | 726.4 | 106.0 | 182.0 | 238.0 | 115.6 | 219.5 | 30.6 | 149.5 | 32.5 | 0.46 | 2305.4 | 0.121 |
| PU | B | 41.8 | 469.5 | 264.2 | 163.4 | 189.4 | 7.9 | 203.2 | 68.9 | 149.5 | 13.9 | 0.64 | 4128.8 | 0.120 |
| PU | C | 576.4 | 1107.0 | 340.4 | 190.2 | 596.6 | 115.9 | 368.1 | 26.4 | 149.5 | 40.7 | 0.87 | 3912.6 | 0.117 |
| PU | D | 167.8 | 584.7 | 247.4 | 169.4 | 242.7 | 42.4 | 244.3 | 55.3 | 149.5 | 19.9 | 0.61 | 4056.2 | 0.119 |
| NU | A | −182.3 | 278.1 | 287.7 | 172.7 | 53.7 | 0[a] | 177.4 | 47.1 | 128.3 | 44.4 | 1.3 | 2213.2 | 0.121 |
| NU | B | 14.8 | 475.8 | 288.8 | 172.1 | 194.7 | 0[a] | 204.4 | 76.8 | 128.3 | 43.8 | 2.2 | 1337.2 | 0.120 |
| NU | C | −61.6 | 410.5 | 297.3 | 174.7 | 180.3 | 0[a] | 186.4 | 43.8 | 128.3 | 46.4 | 1.2 | 2397.6 | 0.119 |
| NU | D | −164.4 | 436.2 | 437.0 | 163.5 | 136.0 | 0[a] | 204.5 | 95.6 | 128.3 | 35.2 | 1.6 | 2731.5 | 0.116 |
| SR | A | −86.8 | 421.1 | 386.0 | 121.9 | 179.5 | 0.23 | 220.8 | 20.8 | 110.2 | 11.7 | 1.5 | 2608.0 | 0.121 |
| SR | B | 45.5 | 588.1 | 229.7 | 312.9 | 226.3 | 144.7 | 152.0 | 65.2 | 110.2 | 202.7 | 1.2 | 1946.3 | 0.120 |
| SR | C | 288.6 | 578.3 | 175.0 | 114.6 | 319.6 | 0.0[b] | 235.4 | 23.3 | 110.2 | 4.4 | 0.61 | 2869.6 | 0.120 |
| SR | D | 139.6 | 630.5 | 361.1 | 129.8 | 397.0 | 101.1 | 102.8 | 29.6 | 110.2 | 19.6 | 2.6 | 1415.9 | 0.120 |
| CG | A | −1206.8 | 89.9 | 422.1 | 874.7 | 32.1 | 0[a] | 44.7[c] | 13.1 | 137.4 | 737.3 | 1.4 | 3126.4 | 0.106 |
| CG | B | −440.3 | 58.2 | 350.3 | 148.3 | 7.0 | 0[a] | 44.7[c] | 6.6 | 137.4 | 10.9 | 1.7 | 2097.4 | 0.107 |
| CG | C | −1335.3 | 60.3 | 827.3 | 568.3 | 0.1 | 0[a] | 41.5 | 18.7 | 137.4 | 430.9 | 3.5 | 2357.1 | 0.121 |
| CG | D | −597.9 | 79.3 | 406.2 | 271.0 | 0.1 | 0[a] | 47.8 | 31.4 | 137.4 | 133.6 | 1.5 | 2655.1 | 0.119 |

Note: Carbon stocks of SOM at 0-10 cm is obtained from Abe et al. (2024).

Superscript a: the $I_u$ in NU and CG was assumed to be zero because these stand types have no understory vegetation or have only small patches of moss species, respectively.

Superscript b: the $I_u$ was assumed to be zero because the understory biomass in 2023 was lower than that in 2022.

Superscript c: the $L$ of the CG was measured on two plots. Therefore, the SD and statistical differences were not evaluated.

**Table S2**. Equations for estimating biomass and $CO_2$ efflux

| Description | Equation | Source |
|---|---|---|
| **Individual-level overstory tree biomass of Asebi (*W*, kg)** | | |
| Stem and branch weight | $\log_{10}(W) = -1.6586 + 2.3696 \log_{10}(D)$ | a |
| Coarse root weight | $W = 37.30 + 0.026D^{2.224}e^{-3.432}$ | b |
| **Individual-level overstory tree biomass except for Asebi (*W*, kg)** | | |
| Stem weight | $\ln(W) = -1.515 + 1.647 \ln(D) + 0.380 \ln(D)^2 - 0.056 \ln(D)^3 + 0.81 \ln(\rho)$ | c |
| Branch weight for deciduous angiosperms | $\ln(W) = -4.314 + 2.502 \ln(D)$ | c |
| Branch weight for evergreen angiosperms | $\ln(W) = -3.964 + 2.400 \ln(D)$ | c |
| Branch weight for evergreen gymnosperms | $\ln(W) = -4.189 + 2.276 \ln(D)$ | c |
| Coarse root weight for deciduous angiosperms | $\ln(W) = -3.274 + 2.315 \ln(D)$ | c |
| Coarse root weight for evergreen angiosperms | $\ln(W) = -3.432 + 2.224 \ln(D)$ | c |
| Coarse root weight for evergreen gymnosperms | $\ln(W) = -3.786 + 2.345 \ln(D)$ | c |
| **Understory vegetation biomass (*B*, g $m^{-2}$)** | | |
| Dwarf bamboo | $B = 15.93NH_{\mathrm{d}} + 335$ | d |
| Other understory vegetation | $B = 2706.3AH_{\mathrm{o}} - 108.40$ | e |
| **$CO_2$ efflux from surface soil except for root respiration (*F*, µmol $CO_2$ $m^{-2}$ $s^{-1}$)** | | |
| For PU plot | $F = 0.096 \exp(0.10T_{\mathrm{s}})$ | d |
| For NU and CG plot | $F = 0.072 \exp(0.11T_{\mathrm{s}})$ | d |
| For SR plot | $F = 0.051 \exp(0.12T_{\mathrm{s}})$ | d |
| **$CO_2$ efflux from coarse woody debris ($R_{\mathrm{h\_CWD}}$, g C $m^{-2}$ $yr^{-1}$)** | | |
| | $R_{\mathrm{h\_CWD}} = M - M \exp(-kt)$, $k = 0.273 + 0.00000668T_{\mathrm{a}} + 0.00000223P_{\mathrm{c}} - 0.00370L_{\mathrm{a}} - 0.0000383A_{\mathrm{l}} + 0.0000678S_{\mathrm{f}}$ | f |

**Abbreviation**: $D$ is diameter at breast height (cm) or diameter at 5 cm position above ground for Asebi (cm), $\rho$ is species-specific wood gravity (g $m^{-3}$), $N$ *is* the number of culms of dwarf bamboo ($m^{-2}$), $H_d$ is the culm height of dwarf bamboo (m), $A$ is the vegetation coverage of understory vegetation excluding dwarf bamboo ($m^{-2}$), $H_o$ is the community height of understory vegetation excluding dwarf bamboo (m) inside the subplot, $T_s$ is soil temperature (°C), $M$ is carbon stock of coarse woody debris (g C $m^{-2}$), $k$ is decay constant (g $g^{-1}$ $yr^{-1}$), $T_a$ is mean annual air temperature (°C), $P_c$ is annual precipitation (mm), $L_a$ is latitude (°N), $A_l$ is altitude (m), and $S_f$ is annual snowfall (kg $m^{-2}$).

**Sources**:

a: Ichihashi, R., Katayama, A., 2024. Aboveground biomass and structural characteristics of poisonous Pieris japonica shrub stands dominating under deer pressure. J. Forest Res. 29, 1-5 (Online early). https://doi.org/10.1080/13416979.2024.2370065

b: Abe, H., Kume, T., Katayama, A., 2024. Reduction in forest carbon stocks by sika deer-induced stand structural alterations. For. Ecol. Manage. 562, 121938. https://doi.org/10.1016/j.foreco.2024.121938

c: Ishihara, M.I., Utsugi, H., Tanouchi, H., Aiba, M., Kurokawa, H., Onoda, Y., Nagano, M., Umehara, T., Ando, M., Miyata, R., Hiura, T., 2015. Efficacy of generic allometric equations for estimating biomass: a test in Japanese natural forests. Ecol. Appl. 25, 1433–1446. https://doi.org/10.1890/14-0175.1

d: Abe, H., Kume, T., Katayama, A., 2025. Soil respiration and heterotrophic respiration in a cool-temperate forest with sika deer-induced alteration of understory vegetation. J. Agric. Meteorol. D-24-00032. https://doi.org/10.2480/agrmet.d-24-00032

e: Fig. S2 in this supplemental materials

f: Dai, Z., Trettin, C.C., Burton, A.J., Jurgensen, M.F., Page-Dumroese, D.S., Forschler, B.T., Schilling, J.S., Lindner, D.L., 2021. Coarse woody debris decomposition assessment tool: Model development and sensitivity analysis. PLoS One 16, e0251893. https://doi.org/10.1371/journal.pone.0251893

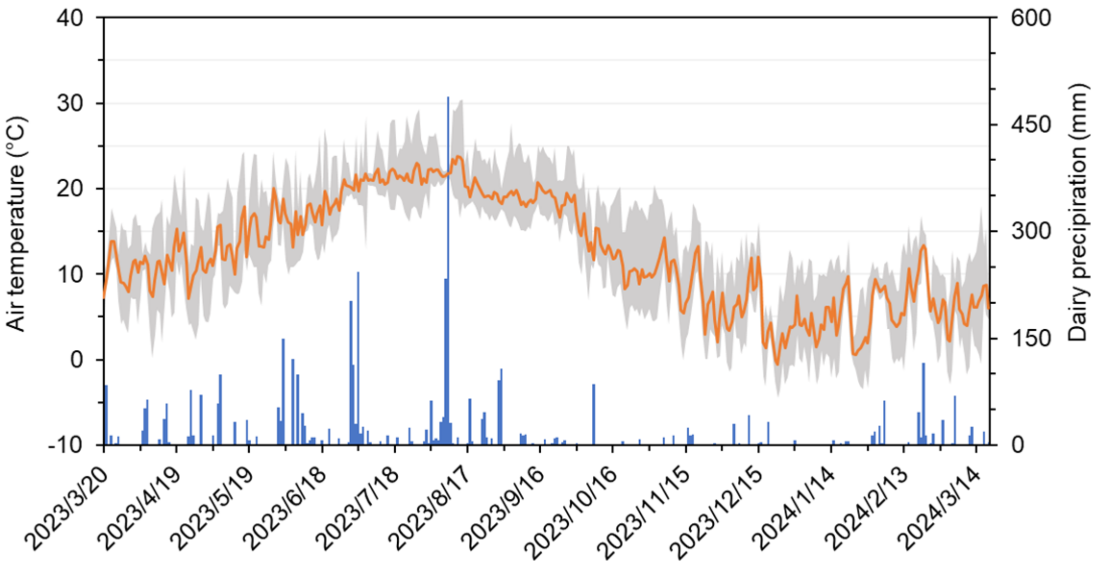


**Fig. S1.** Seasonable variation of air temperature and precipitation in the study area. Orange line indicate daily mean temperature. Shaded area indicates daily maximum and minimum temperature. Blue bars indicate daily total precipitation.

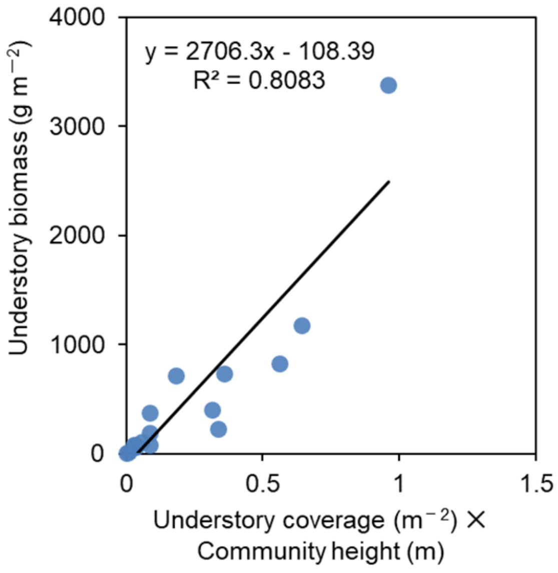


**Fig. S2.** Relationships between understory biomass and multipliers for understory coverage and community height. Data are obtained by Abe et al. (2024c).

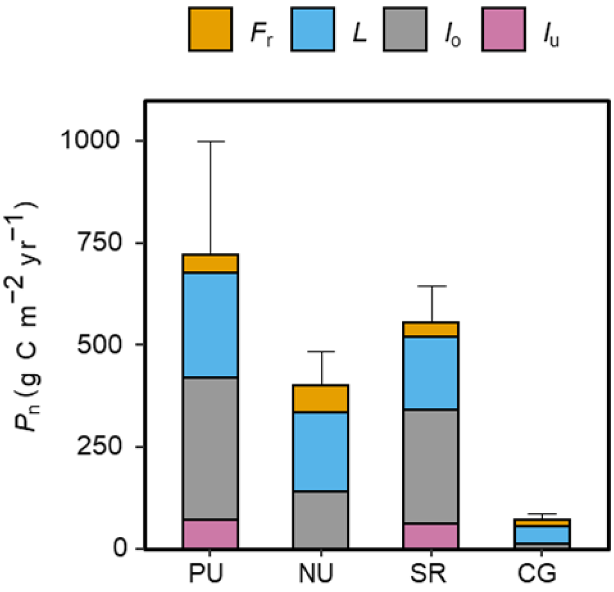


**Fig. S3.** Values of net primary production ($P_n$) components in each stand type. Stacked bars show the mean value of each component. Error bars indicate the SD of total $P_n$. Abbreviations for $P_n$ components: $F_r$; fine root production, $L$; litter production, $I_u$; stand biomass increment of understory vegetation, and $I_o$; stand biomass increment of overstory trees.

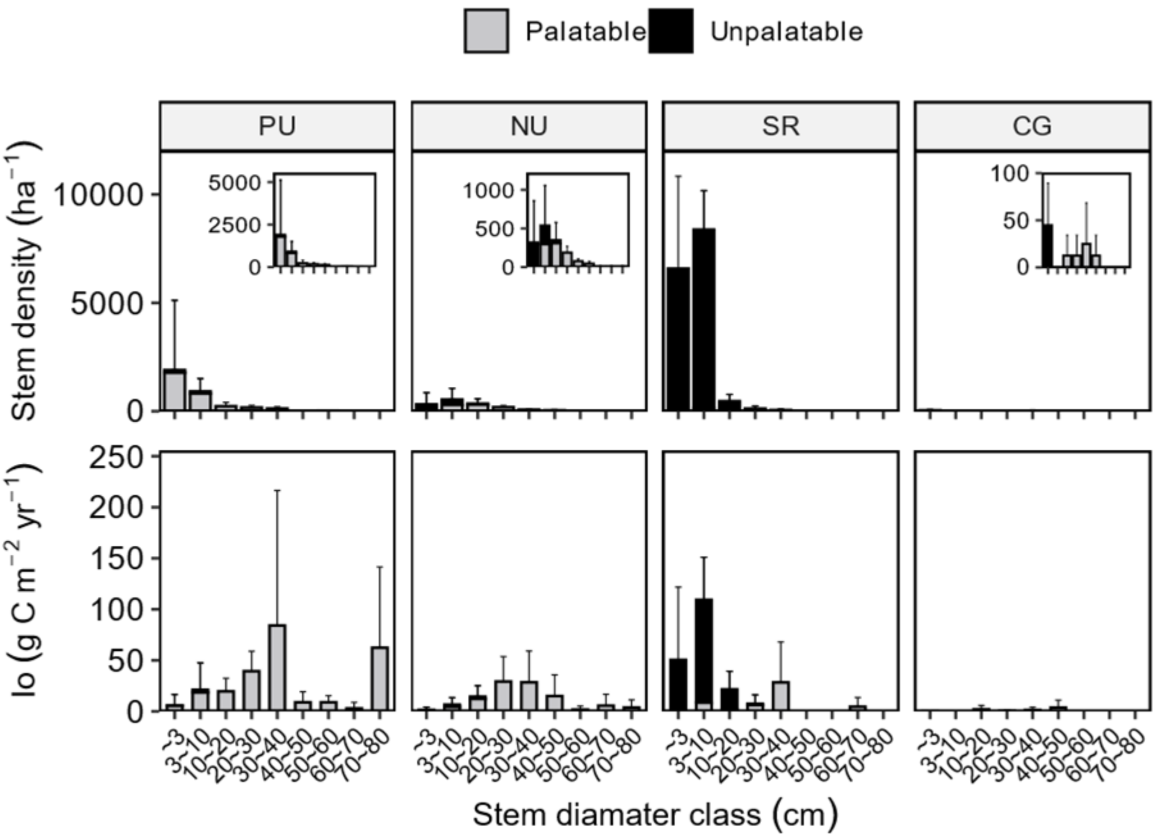


**Fig. S4.** Stem diameter (*D*) class distribution of stem density (top panel) and stand increments of overstory woody biomass ($I_o$) (bottom panel). The stacking bars indicate the mean value for palatable or unpalatable tree species obtained from each plot. Error bars indicate the SD of the total value for each *D* class. Note that *D* includes the diameter at 5 cm height ($D_5$) of Asebi *(Pieris japonica*) and the diameter at breast height (DBH) of other tree species.

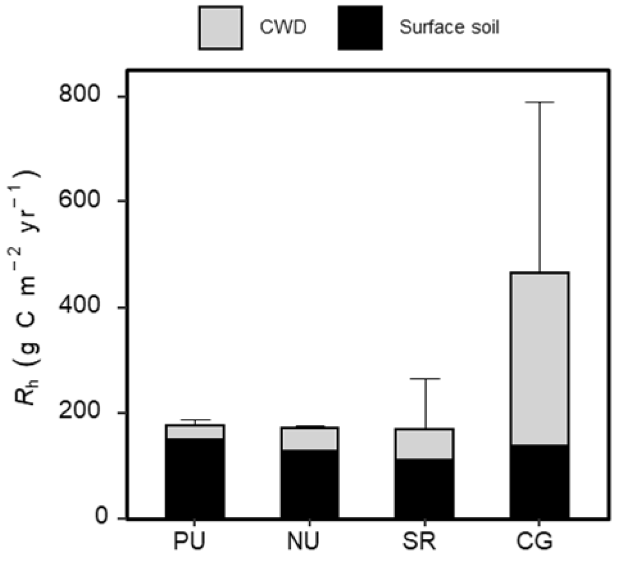


**Fig. S5.** Values of heterotrophic respiration ($R_h$) components in each stand type. Stacked bars show the mean value of each component. Error bars indicate the SD of total $R_h$.

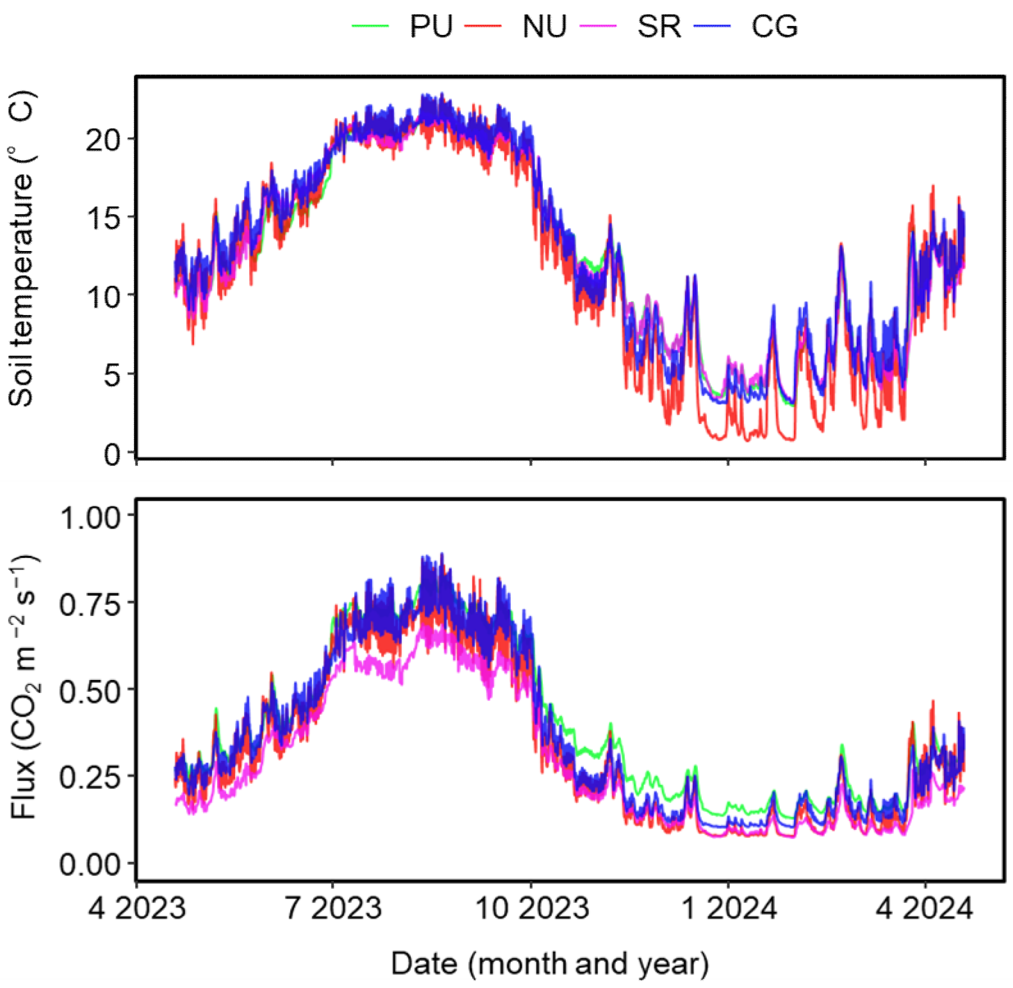


**Fig. S6.** Seasonal trends for soil temperature (upper panel) and $CO_2$ efflux from soil surface without root respiration (bottom).

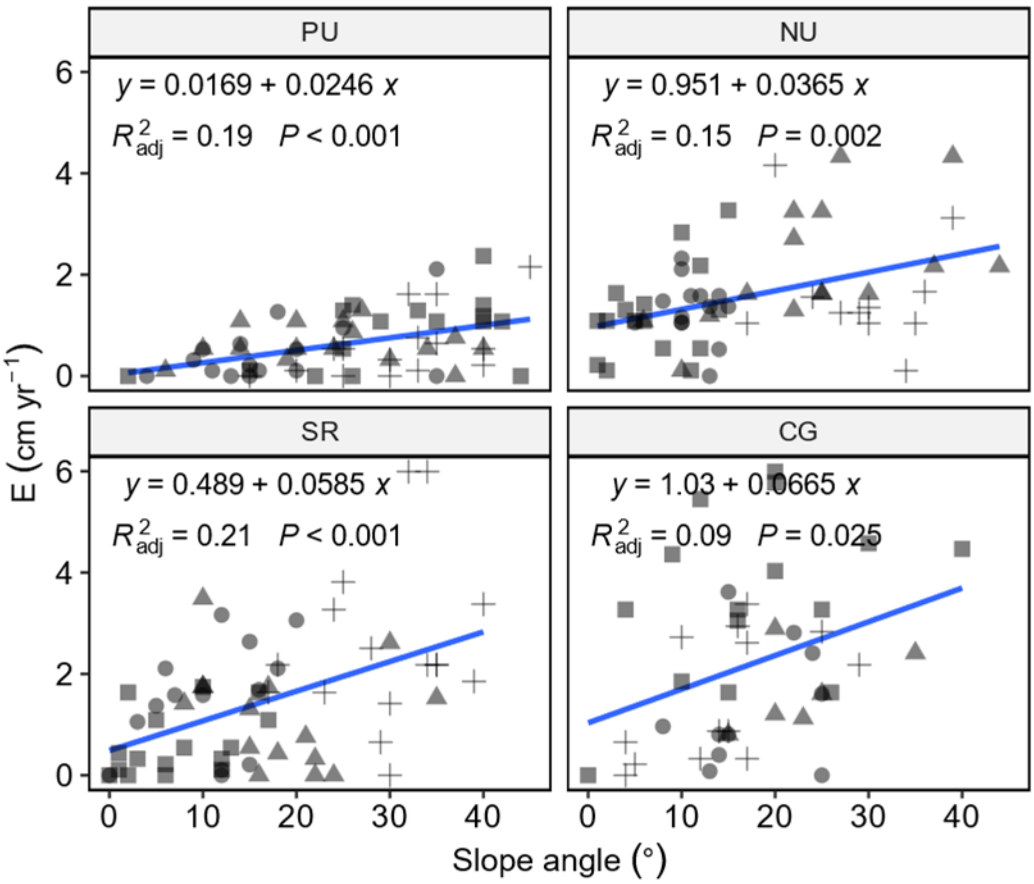


**Fig. S7.** Effect of slope angle (°) on eroded soil depth (*E*) in each stand type. Dots refer to the result of each pins. Symbols (circle, triangle, square, and cross) refer to the plot replicates in each stand type. Blue line indicates the regression line between *E* and slope angle. The regression equation with adjusted $R^2$ and *p*-value were shown inside the graph.
